# Explainable Artificial Intelligence Reveals Potential Candidate Mechanism of Strain-Specific Drug Depletion

**DOI:** 10.64898/2026.01.23.701358

**Authors:** Noorul Fathima Abdul Kafoor, Moe Elbadawi

## Abstract

Oral medications can be bioaccumulated or metabolised by gastrointestinal bacteria in a process collectively termed ‘drug depletion’. The precise biological mechanisms governing strain-specific depletion remain poorly understood, and systematic experimental classification of drug-strain interactions via *in vitro* studies is both costly and time-consuming. In this study, artificial intelligence (AI) methodologies combining machine learning (ML) and natural language processing (NLP) were applied to predict strain-specific drug depletion. The dataset comprised 16,802 drug-strain interaction pairs, with drugs represented by physicochemical descriptors and bacterial strains represented by whole-genome sequences. NLP techniques were used to transform genomic data into feature representations suitable for ML model training. The resulting models achieved strong predictive performance, with a balanced accuracy of 0.90 ± 0.02 and Matthews correlation coefficient of 0.54 ± 0.10. Feature importance analysis revealed that both drug properties and genomic features contributed to model predictions. Among the highest-ranking genomic features, BLASTX annotation identified several enzymes with known or plausible roles in drug metabolism. To further explore the mechanistic relevance of these features, two candidate enzymes were selected for molecular docking against drugs experimentally observed to be depleted. Glycosidase was found to possess binding energies of -8.69 and -7.88 kcal/mol for the two cardiac glycoside drugs digitoxin and digoxin, respectively; whereas acetyl-CoA carboxylase biotin carboxylase presented with binding energies for between -7.09 and -7.74 kcal/mol at one of its druggable sites. Collectively, these findings establish a proof-of-concept AI-driven framework that integrates predictive performance with mechanistic interpretability in the study of drug–microbiome interactions. The broader implications and limitations of applying AI in this context are also discussed. These preliminary findings offer a promising strategy for accelerating drug developments through using AI to rapidly highlight potential drug interactions.

**Graphical Abstract:** 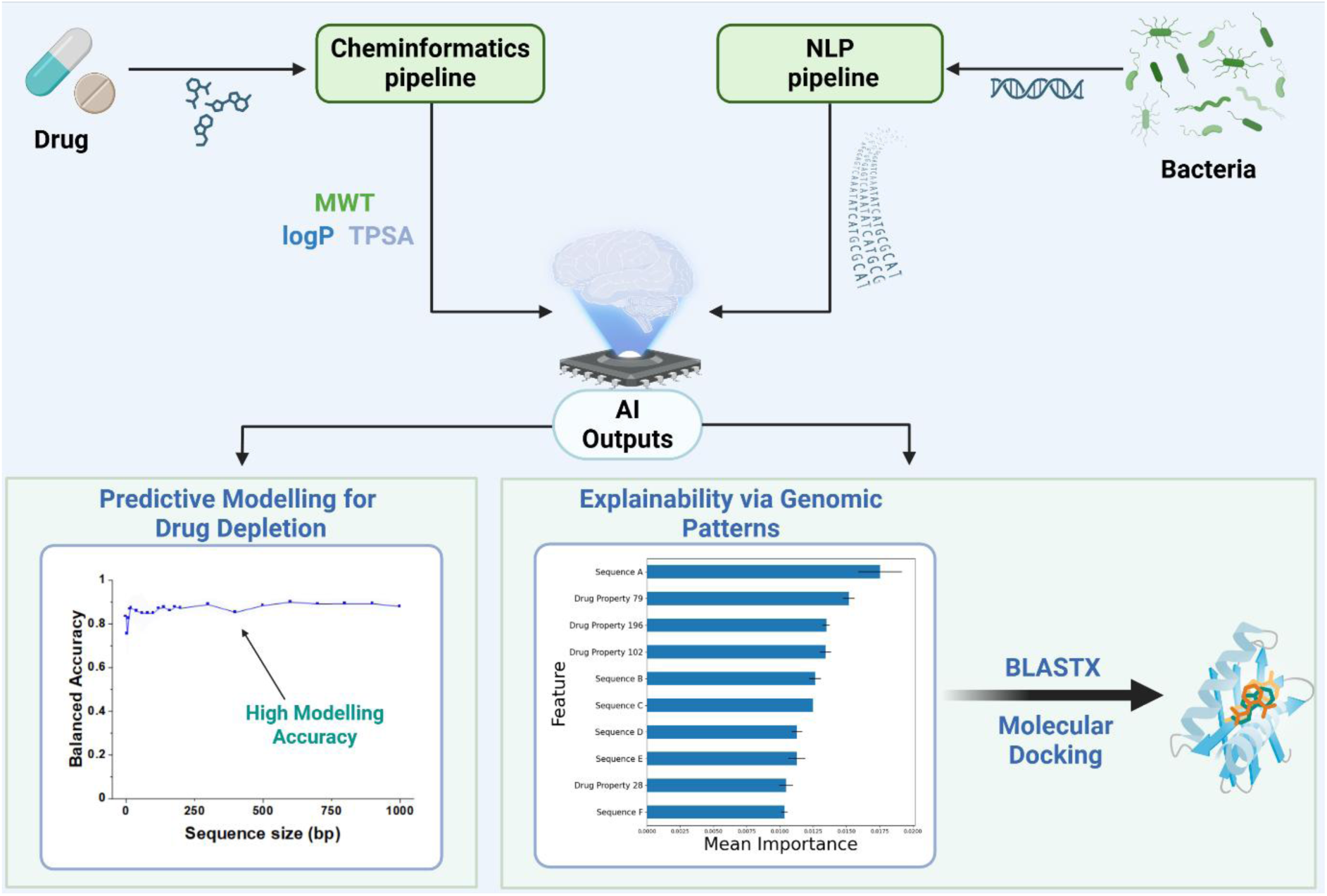

## Introduction

The oral route is the most common form of drug administration, with approximately 60% of small molecules commercially are administered orally [1–4]. It is a preferred route due to the advantages offered, such as patient compliance and non-invasiveness. However, the oral route is also challenging, limiting its preference for all medication. These challenges, which include pH-induced degradation, poor permeability and first-pass metabolism, are widely known in the pharmaceutical field [5–7]. Unfortunately, emerging evidence suggests an additional complication to drug bioavailability by way of the gut microbiome [8].

The human gut microbiome is composed of trillions of microorganisms wielding substantial metabolic potential [9,10]. As well as the digestion of dietary polysaccharides, synthesis of essential vitamins, and regulation of sex hormones, the intestinal microbiota can significantly alter drugs’ pharmacokinetics [11–14]. Research over the last 5 years has revealed that gut bacteria can deplete the luminal concentrations of drugs via two possible mechanisms: biotransformation and bioaccumulation, in a strain-specific manner [8,15,16]. Biotransformation involves the enzymatic modification of a drug’s chemical structure, leading to the formation of a metabolite with different therapeutic and toxicity profiles than the original drug molecule [17,18]. On the other hand, bioaccumulation does not alter drug structure but entails the uptake of unmodified drug molecules by bacteria and intracellular drug storage [19,20]. Both biotransformation and bioaccumulation have the capacity to promote pharmacokinetic variability between and within individuals, potentially complicating dosing regimens in the clinic and in trials [21,22].

Despite the significance of drug-microbiome interactions, they are not routinely screened for during drug development. Biotransformation and bioaccumulation of drugs by gut microbiota can be experimentally measured *in vitro* by incubating drugs with microbiota over a defined period, and periodically assaying remaining drug concentration in incubation media [12,23,24]. Whilst such protocols have been shown to strongly correlate with *in vivo* intestinal drug absorption, they have not been adopted as part of routine analyses by most pharmaceutical companies [25]. Bringing a drug to market is a notoriously expensive process, therefore companies may wish to avoid incorporating additional assays to their pipelines. For this reason, the *in silico* prediction of drug-microbiome interactions presents an opportunity to identify biotransformation and bioaccumulation at an early stage with minimal resource investment [17,26,27].

Several *in silico* methods have been developed to predict drug biotransformation and/or bioaccumulation by gut microbiota [8,19,28–30]. One emerging method is machine learning (ML), which is transforming research in pharmaceutical sciences, from a costly trial-and-error process, to accurately predicting outcomes [31]. ML, a subset of artificial intelligence (AI), has been reported to accurately predict drug-food interactions, drug-excipient interactions and drug stability in biologically-relevant fluids [32–34]. For drug depletion, recent work has demonstrated that ML can accurately predict drug depletion, with the preliminary work indicating the possibility of accelerating workflows in a cost-effective manner [17]. While the preliminary results are promising, ML suffers from several shortcomings, chief among them are its ‘black box’ approach, which hinders scientific understanding [35]. Furthermore, without a mechanistic understanding of the biodepletion process, generalisation cannot be achieved, which ML is known to suffer from. A drawback of previous work is that ML only considered drug properties as inputs to the model, neglecting to represent bacterial strains during model training. One approach to precisely represent bacteria is to use their genomic sequence. The genome of intestinal bacteria is an important determinant of the microorganisms’ capacity to transform and/or accumulate drugs. This can be exemplified by the fact that gut microbiomes of different individuals have varying capacities to deplete drugs, due to varying abundance of microbiota that encode drug metabolising enzymes or membrane transporters [15–17]. As such, there is potential to understand the underlying molecular mechanism behind drug depletion, thereby expanding our understanding of this emerging phenomenon.

To address these limitations, this study investigates whether integrating bacterial genomic information into ML models can improve both the prediction and mechanistic interpretability of drug biodepletion. Whole-genome sequences were used to represent individual bacterial strains, with natural language processing (NLP) feature extraction enabling their incorporation into predictive models. We systematically evaluate the genomic representation strategy using multiple ML techniques.

Specifically, this work aims to (i) determine the extent to which bacterial genomic data improves model performance for predicting drug depletion, and (ii) assess whether genome-informed models can reveal mechanistic insights into the strain-specific processes governing drug biotransformation and bioaccumulation.

## Materials and Methods

### Data Acquisition

Publicly available Genome Collection File (GCF) assemblies of bacterial strains considered by Zimmermann et al., 2019 [8] were collected between 24^th^ and 27^th^ November 2023 and on 1^st^ July 2024, using resources from The National Centre for Biotechnology Information, U.S. National Library of Medicine (NCBI) [36]. The individual interactions of these 66 bacterial strains with 271 therapeutics were obtained from Zimmermann et al [8]. Data cleaning resulted in a final dataset of 62 strains and 271 drugs, forming 16,802 drug-strain pairs. Here, drug depletion is defined as a percentage decrease in drug concentration equal to or greater than the drug’s adaptive fold-change threshold (DAFCT), with statistical significance (p≤0.05). The classification of certain bacterial strains has changed since the 2019 study; for example, *Bacteroides dorei* and *Bacteroides vulgatus* have been re-classified into the *Phocaeicola* genus [37]. However, names have been retained to maintain consistency with original data and subsequent attempts to predict strain-specific drug depletion.

### Hardware

A standard personal computer was used throughout the experimental procedure: Processor: Intel® Core™ i7-1165G7 CPU, 64 GB DDR4 RAM (3200 MHz).

### Data Pre-processing

#### Bacteria Strain Data Featurisation

Bacterial genome featurisation was performed using an in-house NLP pipeline developed, using Python (NLTK v.3.6.2). Generating the *k*-mer features utilised the FASTA representation of the nucleotide sequence, which were manually extracted from NCBI. The NLP pipeline cleaned the FASTA representation, which involved removing spaces and tabs from the data; and all nucleotide text were converted into uppercase. Then, the NLP pipeline converted the cleaned FASTA representation into *k*-mers using the sequential technique which places *k-*mer windows end-to-end to generate consecutive sequences (Figure 1). The last module of the NLP pipeline utilised *CountVectoriser* from *scikit-learn* to vectorise the *k*-mers into features for ML analysis. The 200 most common *k*-mers across all 62 bacteria are selected and their frequencies in each strain are recorded (Table 2).

**Figure 1:**
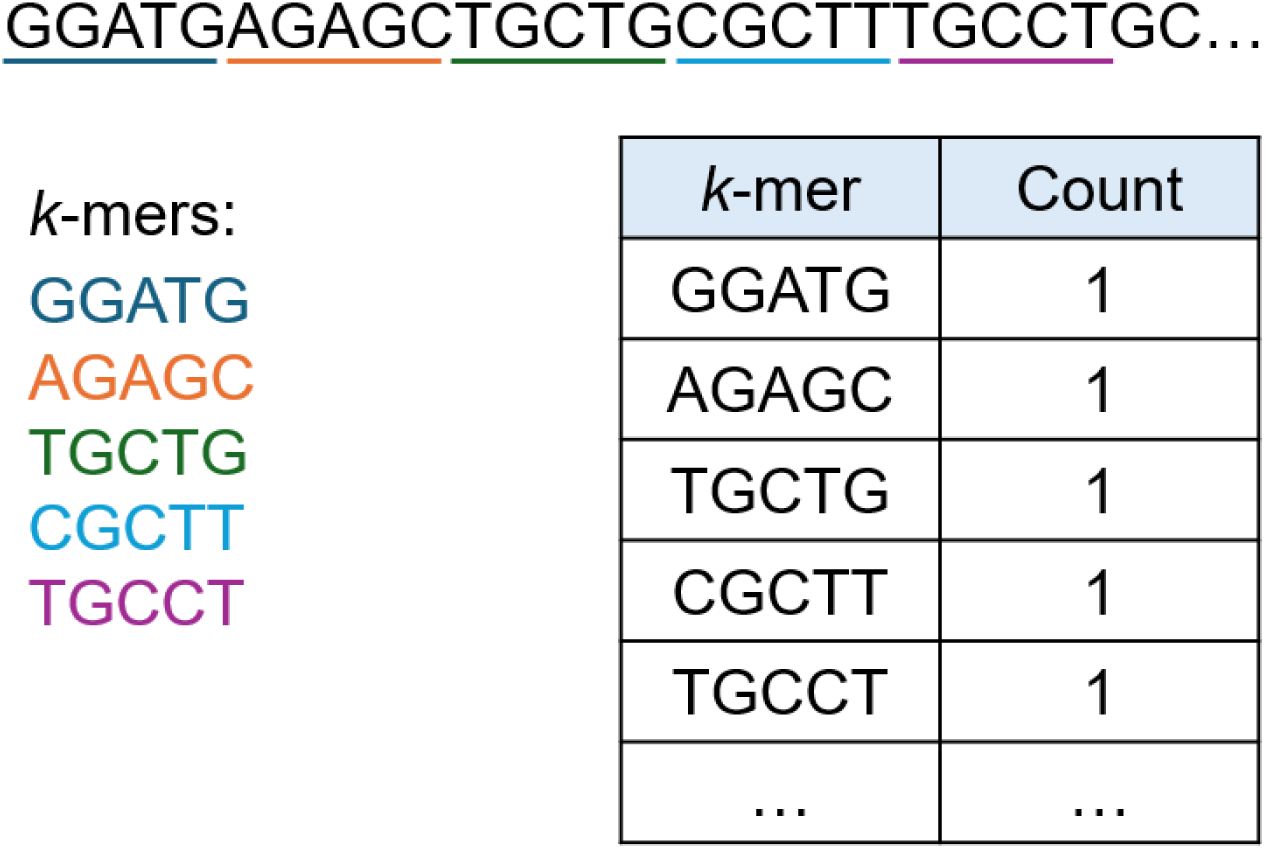
Schematic showing sequential k-mer generation.

**Table 2:** Example k-mer dataset. Dataset structure showing SMILES, drug-strain interaction data, and bacterial features when using the sequential method to generate 6-mers, for each drug-strain pair.

| DrugPair | SMILES | StrainPair | Depletio<br>n | AAAAAA | ... | TTTTTG | TTTTTT |
| --- | --- | --- | --- | --- | --- | --- | --- |
| ABACAVIR<br>SULFATE | C1CC1N<br>C2=C3C(<br>=NC(=N2)<br>N)N(C=N<br>3)C4CC(<br>C=C4)CO<br>.OS(=O)(<br>=O)O | *Akkermansia*<br>*muciniphila*<br>ATCCBAA-835 | No | 366 | ... | 226 | 388 |
| ACEBUTOLO<br>L | CCCC(=O<br>)NC1=CC<br>(=C(C=C1<br>)OCC(CN | *Akkermansia*<br>*muciniphila*<br>ATCCBAA-835 | No | 366 | ... | 226 | 388 |
|  | C(C)C)O) |  |  |  |  |  |  |
|  | C(=O) |  |  |  |  |  |  |
| ... | ... | ... | ... | ... | ... | ... | ... |
| ZOLPIDEM | CC1=CC<br>=C(C=C1)<br>C2=C(N3<br>C=C(C=C<br>C3=N2)C)<br>CC(=O)N(<br>C) | Subdoligranul<br>um variabile<br>DSM15176 | No | 293 | ... | 322 | 284 |

#### Drug Data Featurisation

Simplified Molecular Input Line Entry System (SMILES) representations of therapeutics were used in combination with *RDKit*’s molecular descriptor calculator to generate quantitative structural-properties relationship (QSPR) descriptors for each drug. 8 columns containing missing data were removed from the dataset, resulting in 202 QSPR descriptors (Table 3). These were concatenated with the genomic dataset to form the final dataset following normalisation. For each of the 16,802 drug-strain pairs, 402 input features (200 bacterial and 202 drug features), and one output label (depletion) are given.

**Table 3:** Example QSPR dataset. Dataset structure showing unscaled drug features for each drug-strain pair.

| DrugPair | StrainPair | MaxAbsES<br>tateIndex | MaxEStatel<br>ndex | ... | fr_unbrch_<br>alkane | fr_urea |
| --- | --- | --- | --- | --- | --- | --- |
| ABACAVIR<br>SULFATE | *Akkermansia*<br>*muciniphila*<br>ATCCBAA-835 | 9.25713603<br>472712 | 9.25713603<br>472712 | ... | 0 | 0 |
| ACEBUTOLO<br>L | *Akkermansia*<br>*muciniphila*<br>ATCCBAA-835 | 11.5682245<br>582107 | 11.5682245<br>582107 | ... | 0 | 0 |
| ... | ... | ... | ... | ... | ... | ... |
| ZIPRASIDON<br>E MESYLATE | *Subdoligranulu*<br>*m variabile*<br>DSM15176 | 11.5663360<br>896339 | 11.5663360<br>896339 | ... | 0 | 0 |
| ZOLPIDEM | *Subdoligranulu*<br>*m variabile*<br>DSM15176 | 11.8932170<br>152011 | 11.8932170<br>152011 | ... | 0 | 0 |

### Machine Learning

The machine learning (ML) pipeline was developed in Python (v 3.10.2). The cheminformatics library *RDKit* (v 2024.9.3) *MolecularDescriptorCalculator* class generated features representing drug molecular properties. A total of 8 ML techniques were implemented from the *scikit-learn* library (v 1.6.0): random forest classifier (RF), logistic regression (LR), support vector machine classifier (SVM), gradient boosting classifier (GB), decision tree classifier (DTR), multi-layer perceptron classifier (MLP), k-nearest neighbors classifier (KNN), and extra trees classifier (EXTR). Models were evaluated via 5-fold cross-validation with four metrics: balanced accuracy (Eq 1), F1 score (Eq 2), Matthews correlation coefficient (MCC) (Eq 3), and Brier score (Eq 4). The F1 score was set to evaluate performance in predicting the minority class (depletion), ensuring models prioritise clinically significant cases. All four metrics are considered for each of the 8 models, and the mean and standard deviation of each is recorded.

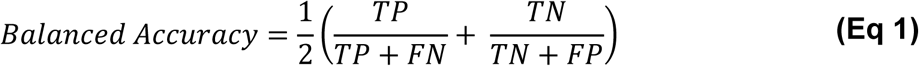

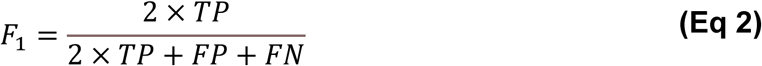

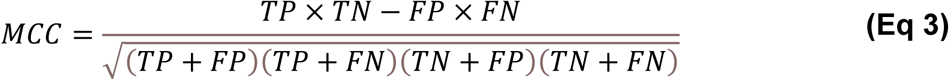

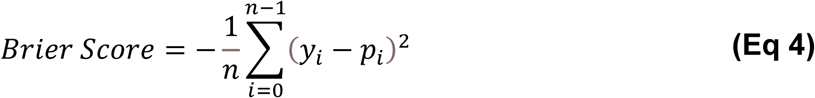

**Key**: **TP** = True Positive, **TN** = True Negative, **FP** = False Positive, **FN** = False Negative, ***n*** = number of samples, ***y*** = actual outcome, ***p*** = predicted probability estimate, ***i*** = iteration.

One-way ANOVA testing of best performers evaluated whether differences in performances are statistically significant (p≤0.05), using the Python package SciPy (v1.16.3). The best-performing model underwent permutation importance analysis to determine the relative importance of features, and the top 25 features were selected for further investigation. Genomic features were analysed using NCBI’s online BLASTX service (nucleotide-to-protein) to identify the encoded proteins [38].

### Molecular Docking

Molecular docking was employed to evaluate the binding of depleted drugs to ML-discovered biological targets, which were glycosidase (PDB ID: 3GM8) and biotin carboxylase (PDB ID: 2VR1). For both PDB structures, all small molecular and non-protein components were removed using ChimeraX (version 1.9) prior to docking. The chemical structures of the depleted drugs were obtained from PubChem in SDF format. Three-dimensional coordinates were generated and hydrogen atoms were added using OpenBabel (version 3.1.1). The structures were then converted to MOL2 format in ChimeraX. Docking calculations were performed using SwissDock with Autodock Vina (SwissDock 2023 release). For glycosidase, the grid size was 20 – 50 - 20 and the box coordinates were 26 - 121 - 82. For biotin carboxylase, the grid size was 50 - 20 - 20 and the box coordinates were 20 - -22 - 23 The docking results were then exported and visualised using ChimeraX.

## Results

### Exploratory Data Analysis (EDA)

Of the 76 bacterial strains investigated by Zimmermann et al., whole genome assemblies were available for 66 strains [8]. Analysis of the available 66 GCFs revealed that 4 strains possessed a completeness value less than 90%, signalling missing genetic data. Hence, these drug-strain pairs were removed from the dataset. In total, 62 bacterial strains are investigated (Figure 2.C), lowering the number of drug-strain pairs explored to 16,802. This data cleaning does not greatly affect depleted pairs but considerably reduces not-depleted pair numbers, improving dataset balance. The final dataset remains imbalanced, with the minority class (depletion) accounting for 13% of the dataset (Figure 2.B).

**Figure 2:**
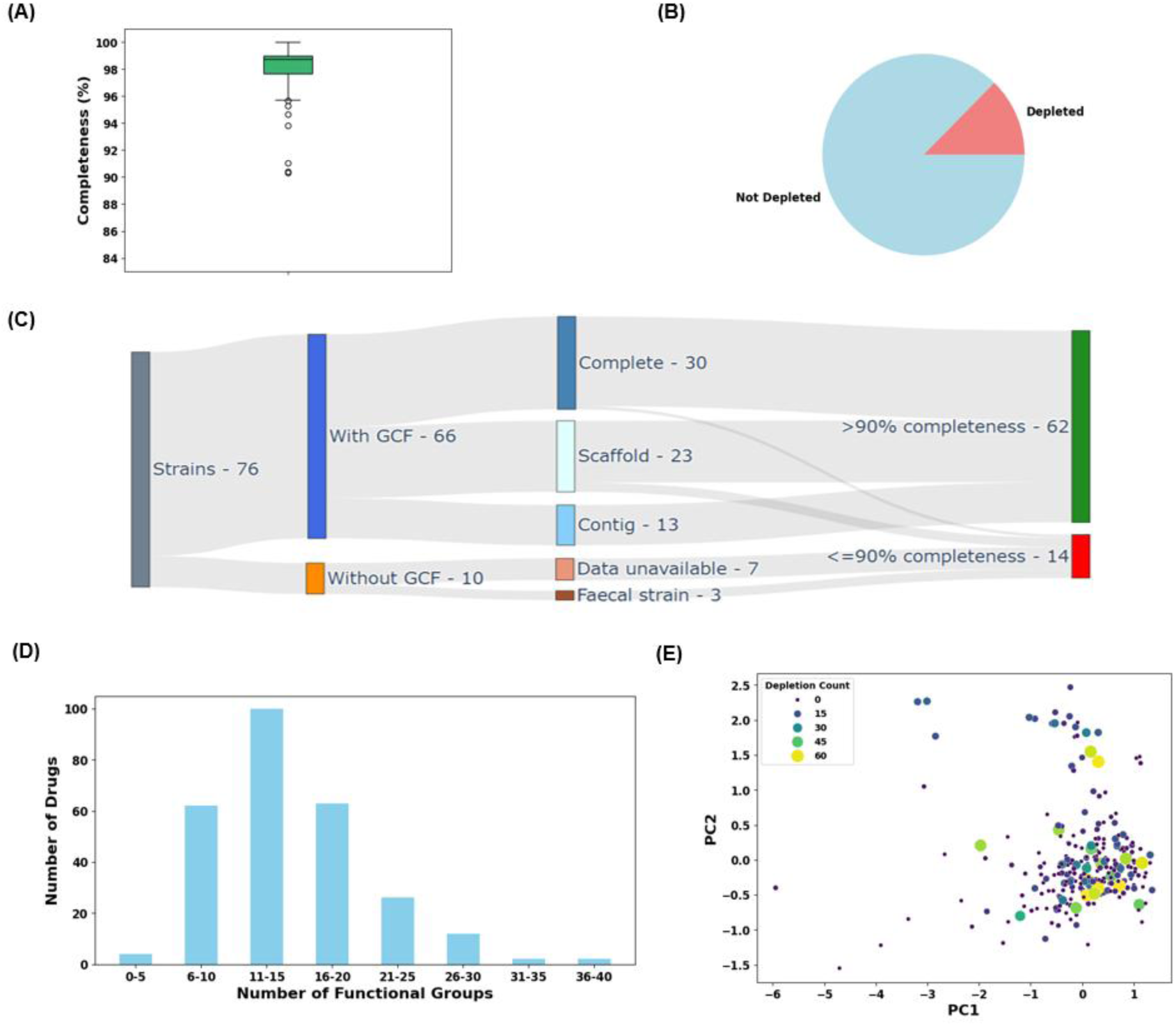
Results of exploratory data analysis. (A) Box plot showing the distribution of completeness percentages of the 66 available bacterial genomes. (B) Pie chart showing that investigation of 62 bacterial strains with 271 therapeutics produces an imbalanced dataset. (C) Sankey diagram showing whole genome sequence availability of bacterial strains and selection of 62 bacterial strains. (D) Bar chart showing the number of functional groups of drugs investigated. (E) Two-dimensional PCA plot, showing the relative chemical space for the 271 therapeutics. Both the colour and sizes of datapoints represent the depletion count.

Figure 2.D demonstrates the varying complexities of the drugs investigated, via the number of functional groups. Principal component analysis (PCA) plot of drug physicochemical properties coloured by depletion count for each drug revealed no obvious clustering between drugs that deplete against those that do not, signalling a complex, non-linear interaction between drug structure and depletion.

A heatmap displaying drug-strain interactions was generated to visualise trends in depletion (Figures 3-5). Here, bacteria and drugs are ordered alphabetically, grouping bacteria by genus and species in the expectation that evolutionarily similar bacteria share drug depletion mechanisms, and therefore display similar depletion patterns. This found bacterial species from the *Bacteroides* and *Parabacteroides* genera exhibited similar depletion patterns, underscoring the benefits of genomic data for understanding depletion. This heatmap also illustrates the variability in number of bacteria depleting drug compounds. Some therapeutics (like abacavir sulfate) are not depleted by any of the considered bacterial strains, while some are depleted by all or most bacterial strains investigated (bisacodyl and artemisinin are depleted by 62 and 61 strains, respectively). This highlights drug depletion is drug-specific and strain-specific, and likely has notable links to drug structural properties and genetic profiles of strains.

**Figure 3:**
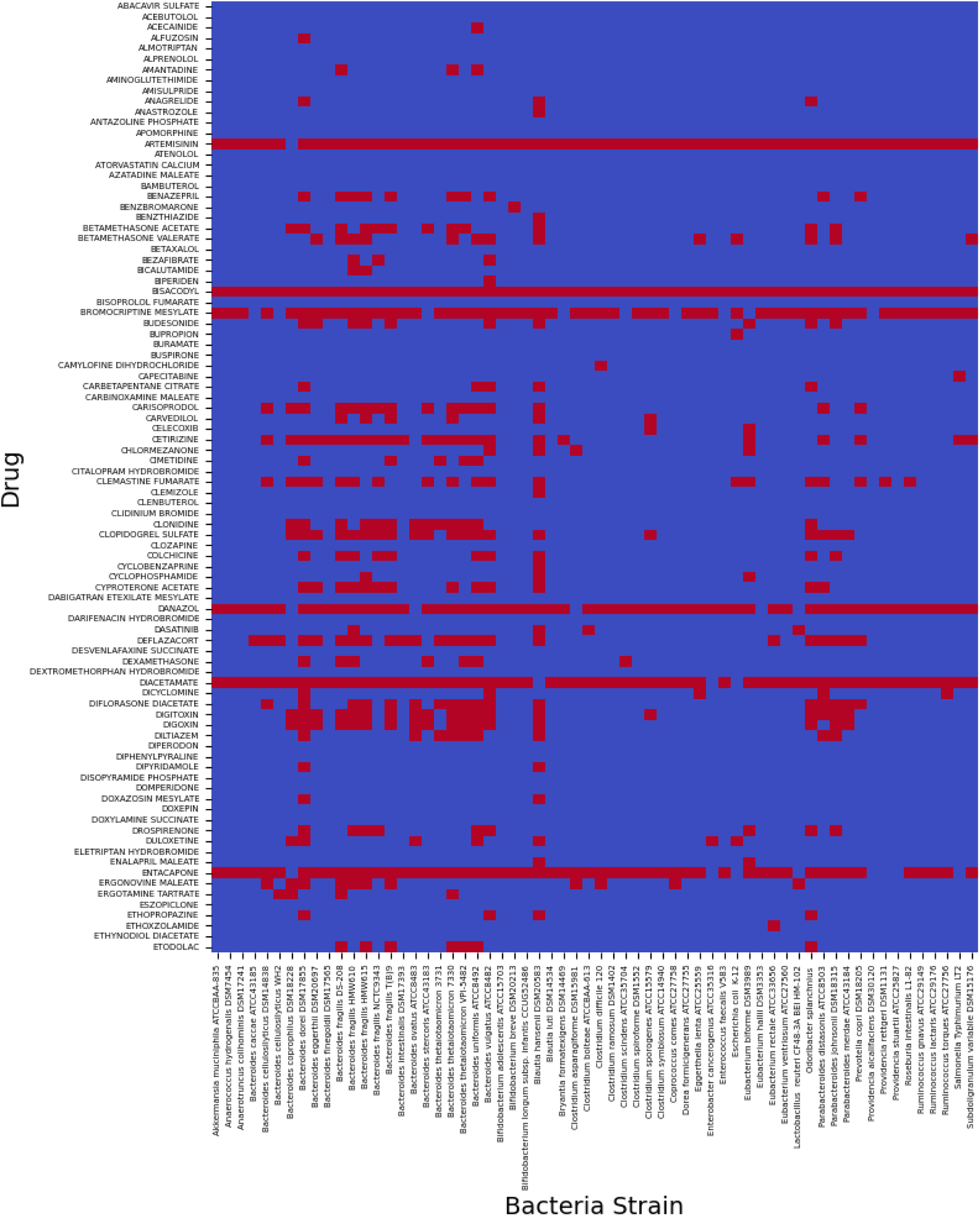
Heatmap showing drug-strain interactions specific to bacterial strains and 90 drugs not shown in figures 4 or 5. Depletion is represented in red, while no depletion is shown in blue.

**Figure 4:**
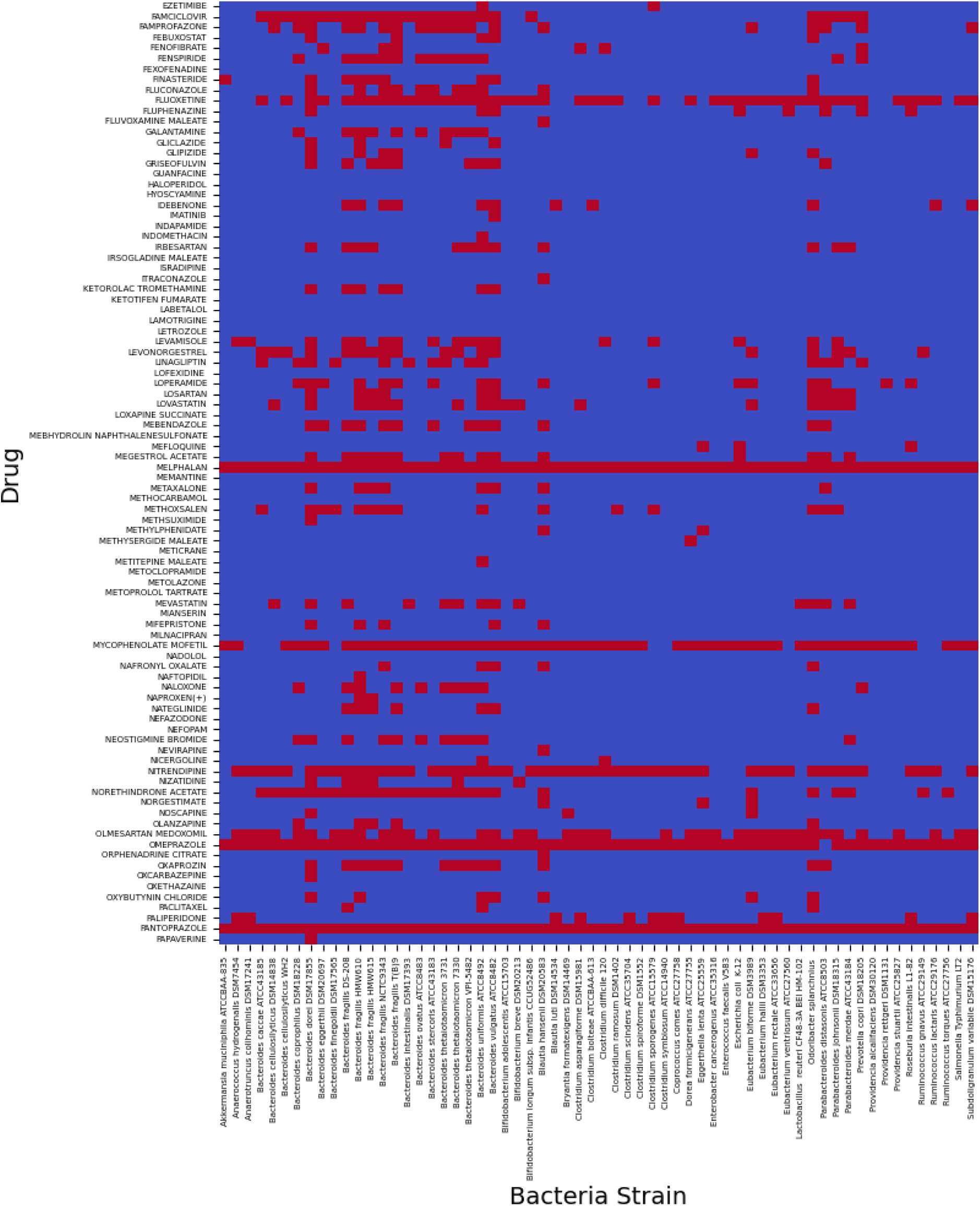
Heatmap showing drug-strain interactions specific to bacterial strains and 90 drugs not shown in figures 3 or 5. Depletion is represented in red, while no depletion is shown in blue.

**Figure 5:**
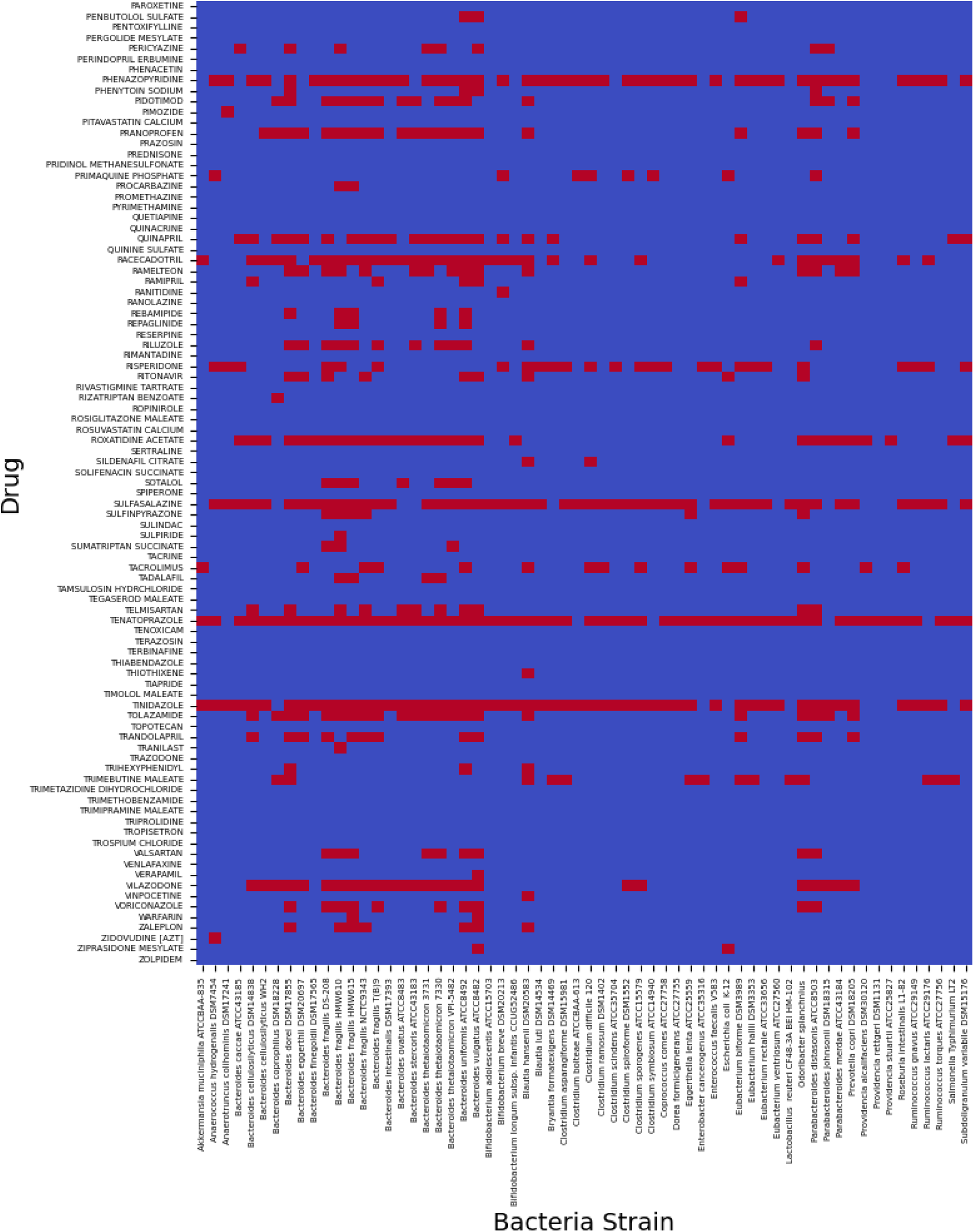
Heatmap showing drug-strain interactions specific to bacterial strains and 91 drugs not shown in figures 3 or 4. Depletion is represented in red, while no depletion is shown in blue.

### Unsupervised Machine Learning

Heatmaps generated from the properties of bacteria and drugs separately have been labelled by depletion count - the number of drugs a strain has been shown to deplete, or the number of strains a drug has been shown to be depleted by (Figures 6 and 7). The hierarchical clustering using the NLP-derived *k-*mer approach revealed bacterial clustering by similar species, confirming that this approach captured the nuances between the different strains (Figure 6). From this multi-variate analysis, there was no obvious correlation between the *k*-mers and depletion count.

**Figure 6:**
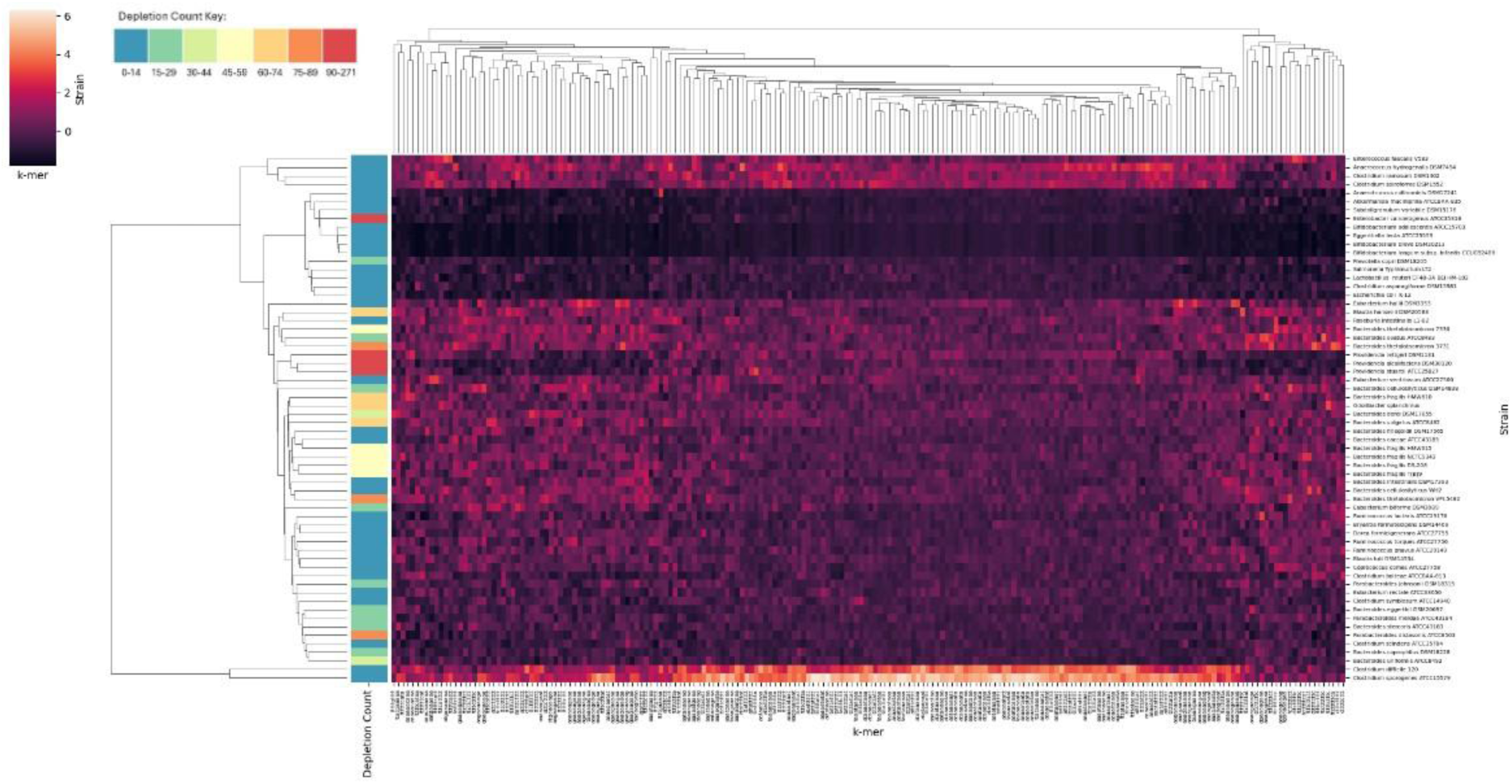
Heatmap showing hierarchal clustering grouping of bacteria by 10-mers, with strains’ depletion categories. Hierarchical clustering is shown through dendrograms.

**Figure 7:**
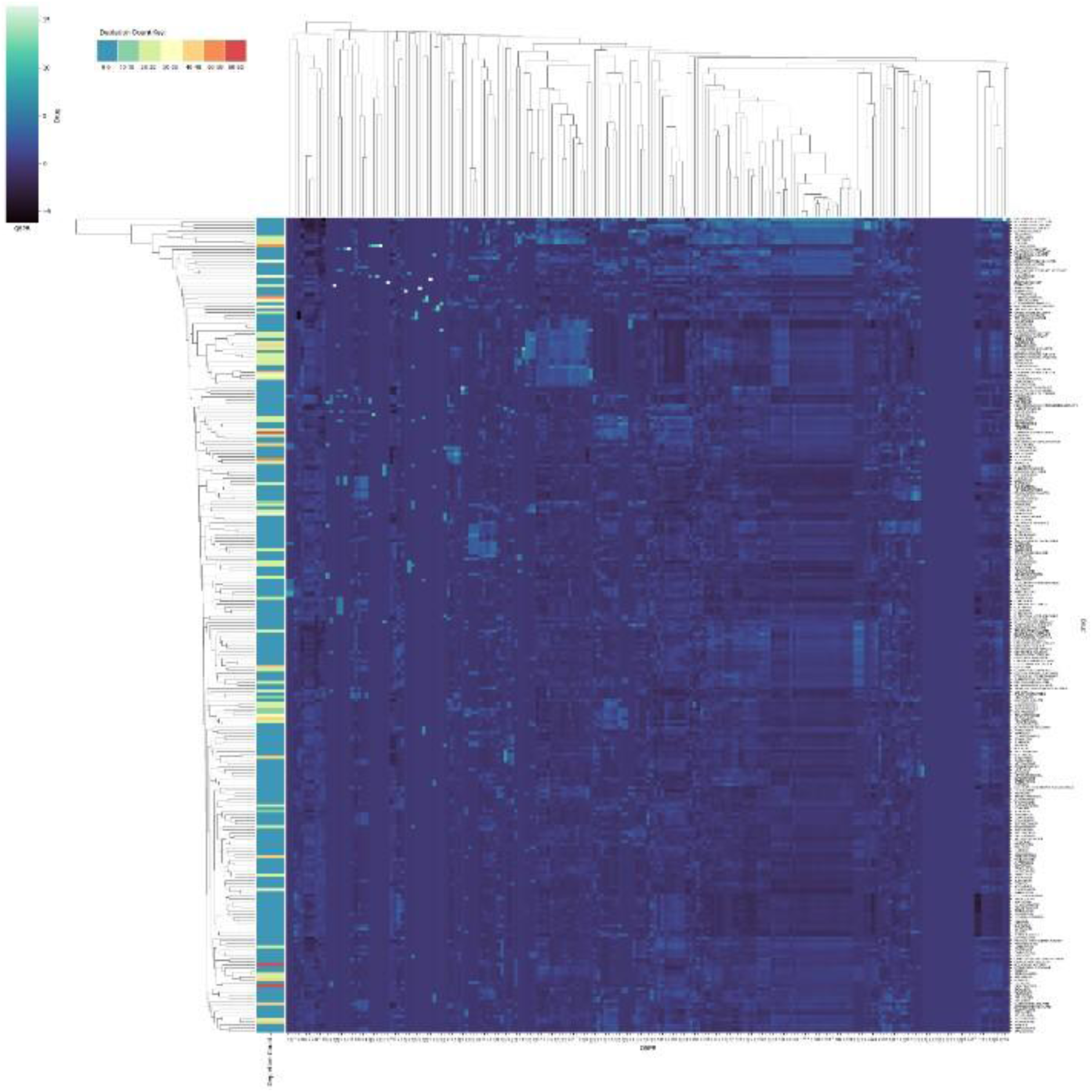
Heatmap showing hierarchal clustering grouping of drugs by physicochemical properties, with drug’s depletion categories. Hierarchical clustering is shown through dendrograms.

The multi-variate analysis of drug depletion using their physicochemical structures were also analysed and similarly, yielded no obvious patterns (Figure 7). These analyses confirm the need for supervised learning to discern a difference between drug-strain pairs that deplete and those that do not.

### Supervised Machine Learning

The effects of differing ML techniques and *k*-mer sizes on model performance were then investigated (Figure 8). Here, a *k* value of zero represents the absence of genetic information. The *k*-mer sizes investigated increase in increments of 5 up to 20; then by 20 until 200; then by 100 up to 1000. Three metrics of model performance (F1 score, balanced accuracy, and MCC), and one representing confidence (Brier score) were investigated. Initial values of all performance and confidence metrics were high across all 8 ML techniques (e.g., balanced accuracies ranged between 82-91%).

**Figure 8:**
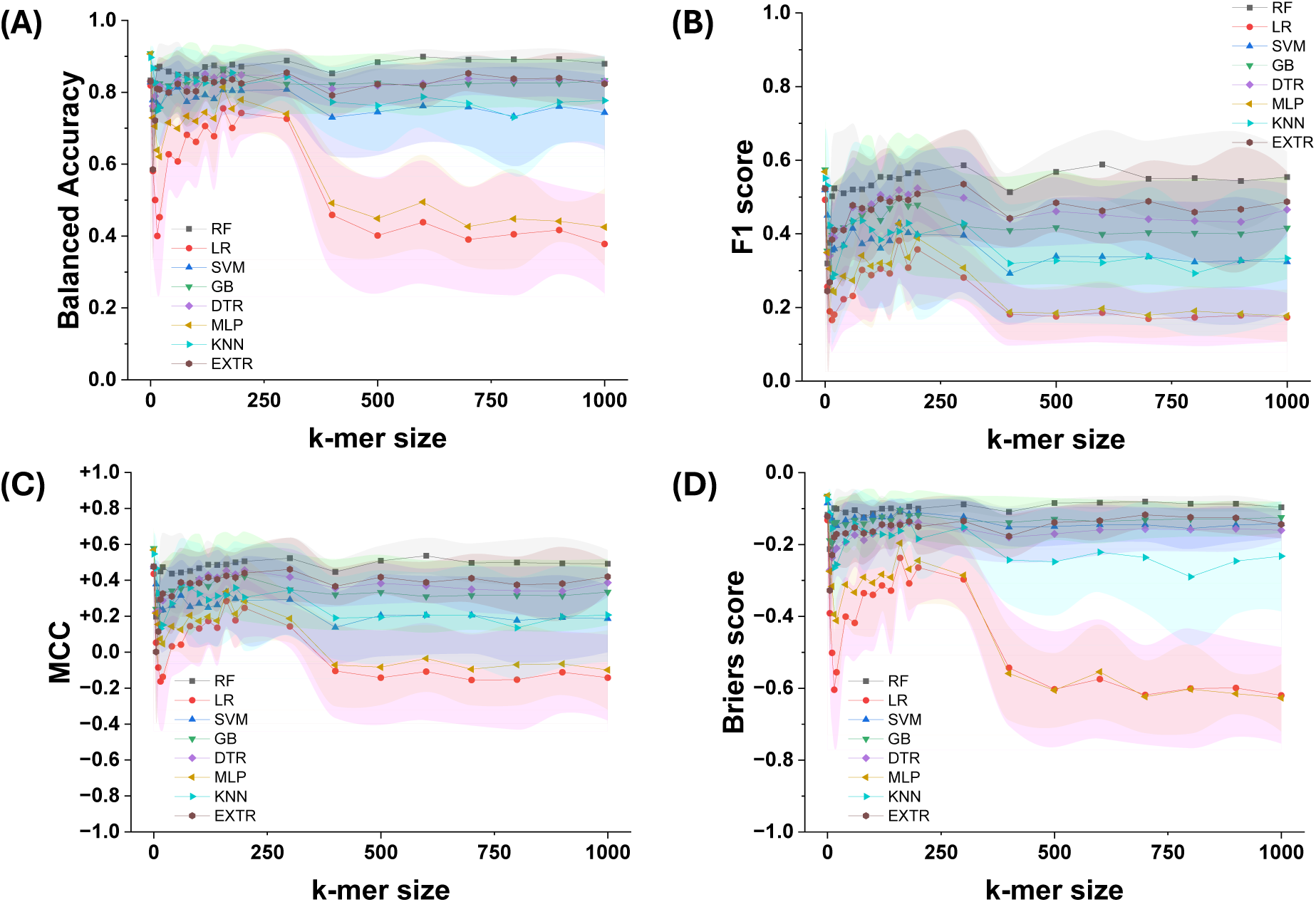
Results of varying ML classifiers and sizes of k-mers. Mean (A) balanced accuracy scores, (B) F1 scores, (C) MCC, and (D) Brier’s scores, as well as relevant standard deviations (shown by shaded region) showing model performance and confidence.

Thereafter, performance decreased dramatically before increasing with *k*-mer size. At 300-mers, LR and MLP performance sharply decreased, whereas the remaining techniques maintained their high performance. In particular, all four metrics suggest tree-based models were the best learners when provided genomic data. For example, an RF model with 600-mers achieved a balanced accuracy, F1 score, MCC and Briers score of 0.90 ± 0.02, 0.59 ± 0.09, 0.54 ± 0.10 and 0.08 ± 0.02, respectively. However, ANOVA revealed no statistical significance between ML techniques using genomic data to that using drug-only properties (i.e., 0-mer), confirming that the addition of genomic data did not degrade ML technique performance through, for example, the addition of noise. Overall, these results revealed that ML techniques yielded a high score, with the choice of *k*-mer and ML technique impacting predictive performance.

### Explainable Machine Learning

Tree-based models can be queried post-prediction to determine which features were important in achieving their prediction. Feature importance analysis was conducted on the top-performing model based on F1 score (RF with 600-mers) and revealed that of the 25 features most important for accurate decision-making, 13 were genomic features (Figure 9). This suggests drug depletion is strain-specific and validates the use of genomic information for prediction.

**Figure 9:**
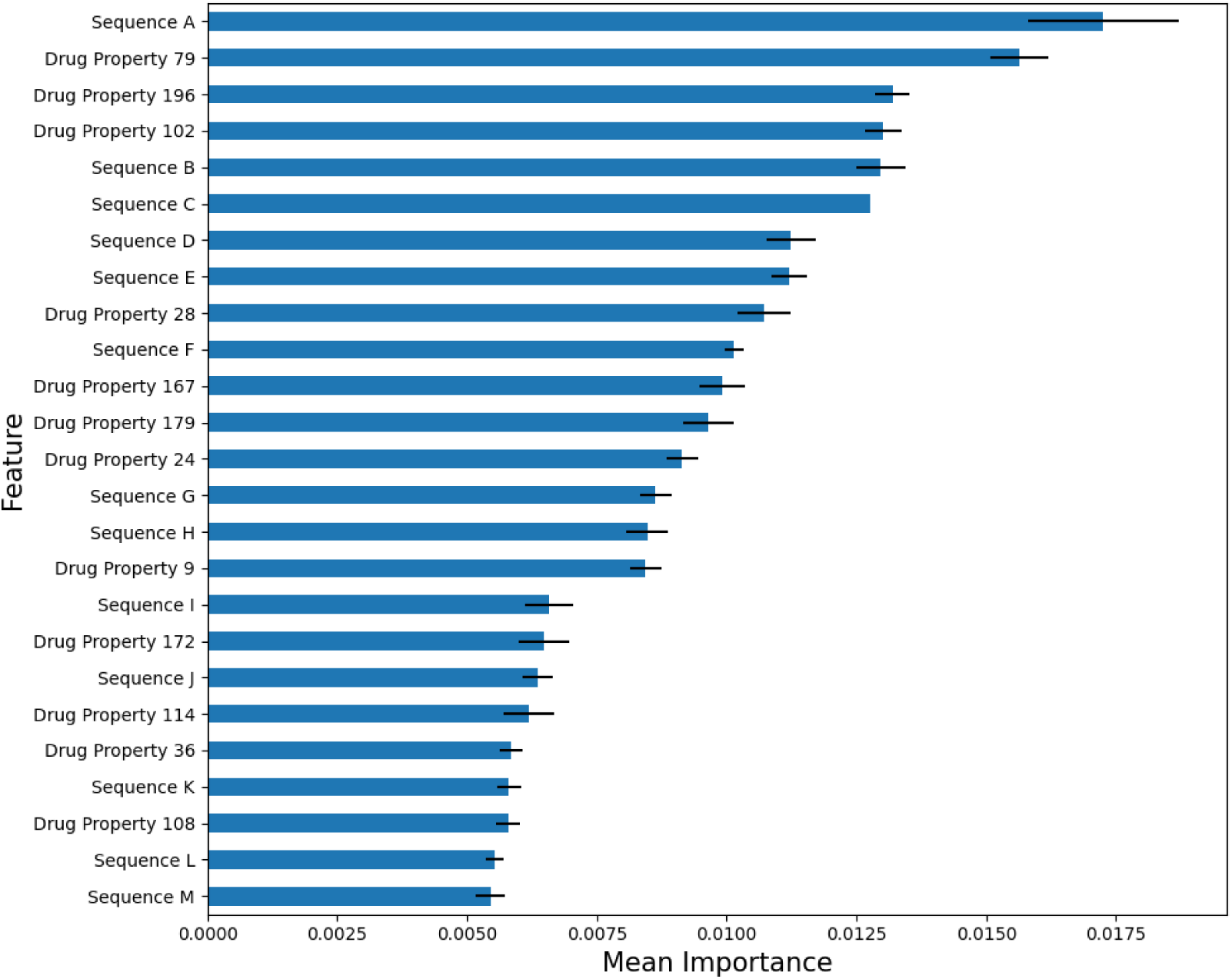
Results of feature importance analysis of model using RF classifier with 600-mers. Refer to table S1 for definitions of drug property numbers.

The sequences identified through feature importance analysis were processed using BLASTX to elucidate whether they pertain to a protein. Table 4 lists the results. The most important feature was sequence A, where BLASTX illuminated that it encoded for glycoside hydrolase enzyme, with both 100% query cover and 100% identity. Glycoside hydrolase is a catalytic enzyme that hydrolyses glycosidic bonds, which are found in compounds included in the training data such as the cardiac glycosides digoxin and digitoxin. Sequence A was only present in *B. vulgatus* ATCC8482, where the strain depleted 31% of 271 drugs investigated. Preliminary molecular docking results revealed that these drugs presented with a relatively high binding energy (Figure 10).

**Figure 10:**
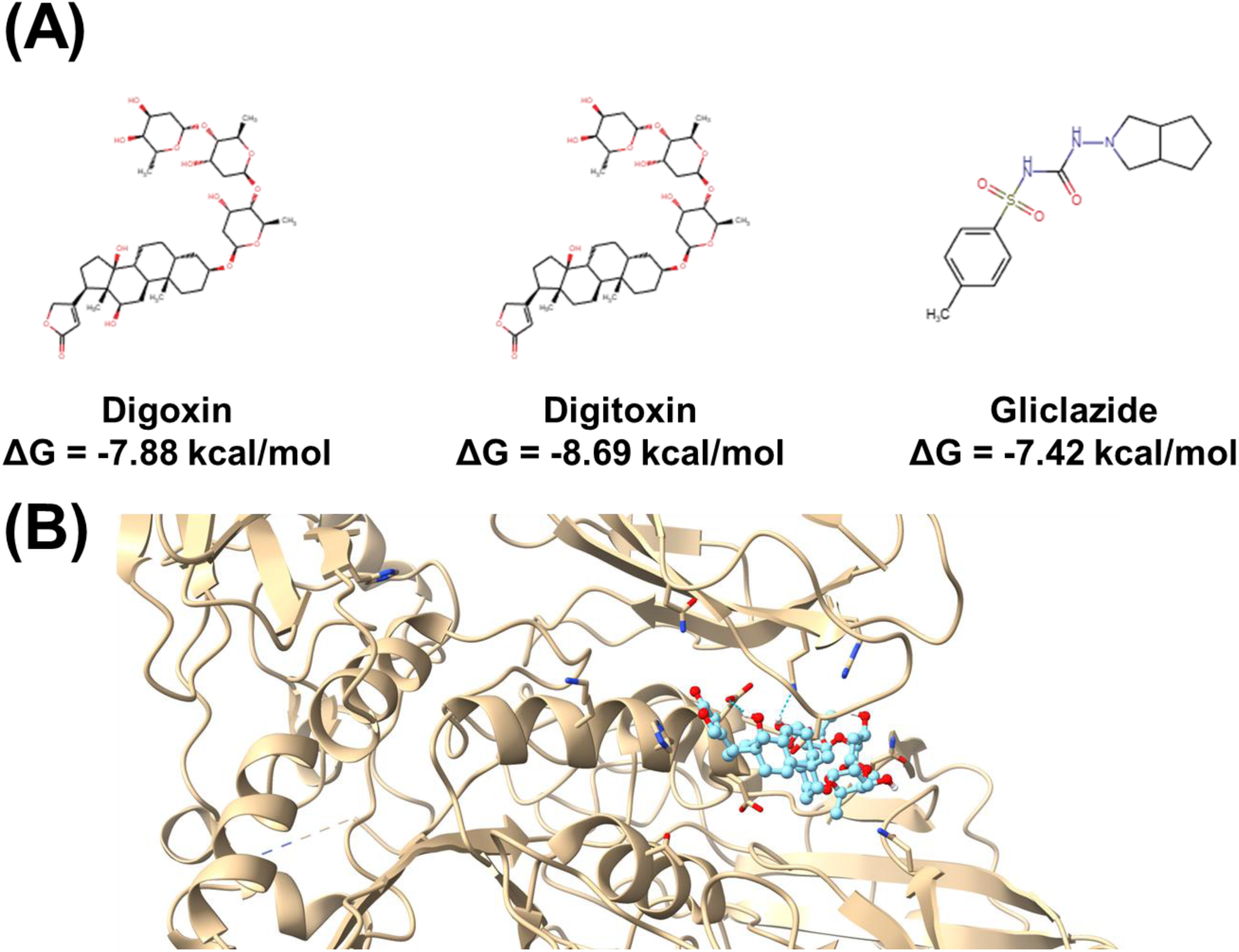
(A) Three compounds depleted by strains containing glycosidase and their corresponding binding affinities. (B) Computational docking of digitoxin as an example demonstrating the potential of the drug binding to a putative binding pocket.

**Table 4:**
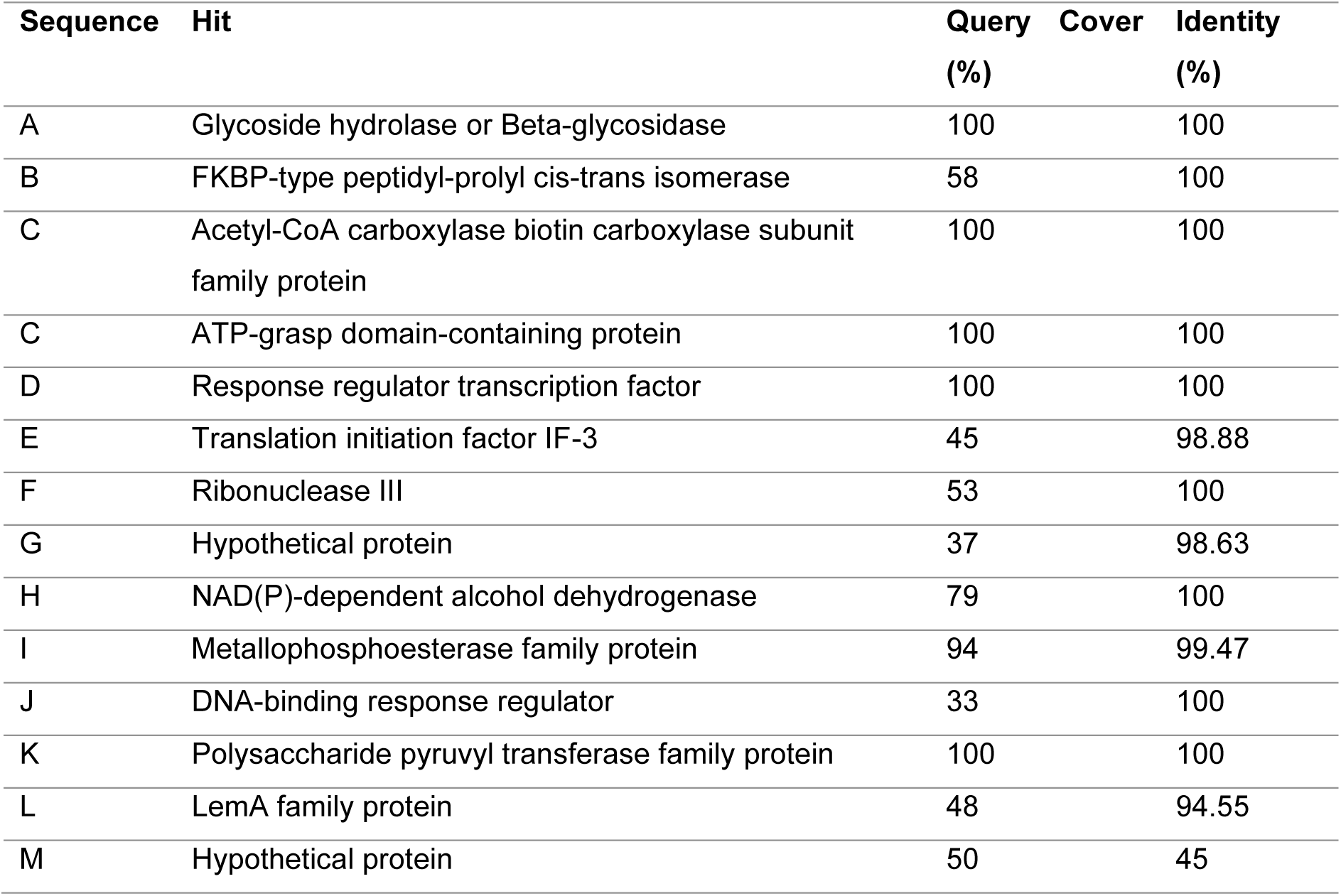
Results of BLASTX searching of genomic sequences identified by feature importance analysis of model using RF classifier with 600-mers.

Another enzyme revealed through BLASTX was acetyl-CoA carboxylase biotin carboxylase subunit family protein, which also exhibited 100% for both query cover and identity. This enzyme’s role is to catalyse the ATP-dependent carboxylation of biotin in the first committed step of fatty-acid biosynthesis. The enzyme has been a subject of drug targeting and thus is known to interact with drugs [39–43]. One region that has been identified as potentially druggable was the ATP-binding site, in particular the adenine-binding pocket. Here, drugs found to be depleted by strains containing the enzyme exhibited moderate binding affinities of approximately -7 kcal/mol (Figure 11). Further experimental validation will be needed to determine whether the enzyme specifically interacts with these compounds or acts as a non-productive off-target binding sink.

**Figure 11:**
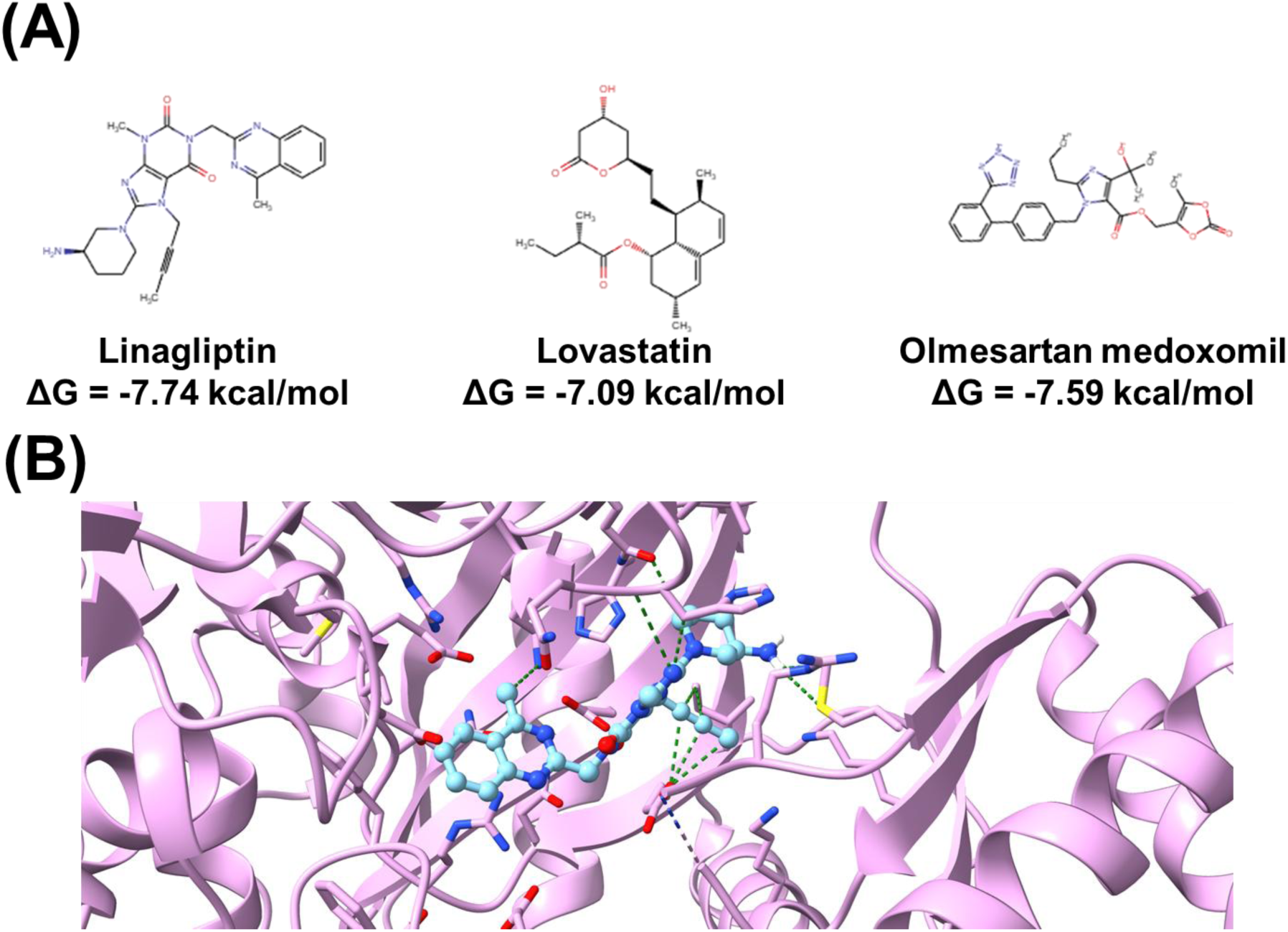
(A) Three compounds depleted by strains containing biotin carboxylase and their corresponding binding affinities. (B) Computational docking of linagliptin as an example demonstrating the potential of the drug binding to the adenine binding pocket.

## Discussion

AI has proven to be a powerful tool in pharmaceutics, and this study lays the groundwork for its use beyond prediction and its potential in enhancing mechanistic understanding of drug depletion [9,44]. The initial study by Zimmerman et al. (2019) proffered some suggestions as to which drug properties were susceptible to depletion, but made no mention of the mechanism of action [8]. This can be attributed to the fact that depletion is a nascent discovery, the complexity of which has been exemplified by EDA (Figure 2). Building upon Zimmermann et al.’s discovery of a causal link between bacterial enzymes and drug depletion, research herein seeks to link genes to mechanisms of depletion, thereby associating them with properties of drugs [8]. Here, we demonstrated that ML is capable of prioritising biologically relevant targets and consequently offering novel insight into mechanisms; namely, glycoside hydrolase-and acetyl-CoA carboxylase-mediated depletion. In doing so, this approach enables ML generalisation to novel, untested drug compounds and bacterial strains, thereby reducing costs and resource expenditure associated with *in vitro* studies.

Our findings establish that both the choice of genomic data and ML technique employed has considerable effects on predictive performance and confidence. LR and MLP classifiers, as well as small *k*-mer sequences resulted in markedly reduced performance. Despite this, no single model consistently outperforms across all metrics. Greatest mean F1 score was found to be in RF with 600-mers, and was taken as the overall best performer. Of models with genomic data developed herein, the 4 best-performing ML techniques were found to be tree-based, which are better suited for classification tasks due to their ability to effectively handle large non-linear datasets [45]. This finding is in line with that of McCoubrey et al., which found the best-performing model to use an EXTR classifier [28]. In the present study, optimal models substantially out-performed previous models (69% balanced accuracy), achieving balanced accuracy scores of 90-91% [28].

Endeavours to reduce the ‘black box’ nature of AI and understand reasoning behind models found that glycoside hydrolase-encoding strain (*B. vulgatus* ATCC8482) depleted both cardiac glycosides investigated (digoxin and digitoxin). These drugs were found to bind a putative binding site on glycosidase with high binding affinity. Glycoside hydrolases cleave glycosidic bonds through hydrolysis reactions [46–48] Thus, this study proposes glycoside hydrolases degrade cardiac glycosides via hydrolytic cleavage of the glycosidic bond connecting the saccharide moiety and steroidal nucleus. This suggests a novel mechanism of bacterial drug depletion, warranting experimental validation.

Another possible explanation is that glycoside hydrolase-mediated depletion may follow lactone ring-targeting pathways, similar to the cardiac glycoside reductase (CGR) enzyme in *Eggerthella lenta* DSM2243 (also known as the ATCC25559 strain) [49]. The number of cyclic esters (lactones) was highlighted by feature importance analysis. Of the 9 drugs containing lactone rings in this study, 3 were not depleted by *B. vulgatus* ATCC8482. Further, Zimmermann et al. did not find *E. lenta* ATCC25559 to deplete either cardiac glycoside [8]. These findings suggest lactone rings alone are not sufficient for cardiac glycoside depletion, with gene expression variations potentially explaining differences in depletion. This is supported by previous studies, which also point towards non-linear relationships between depletion and features [8,50–53]. Therefore, future work may use gene expression profiles in leu of genomic data to reveal active metabolic pathways, allowing greater insight into depletion mechanisms [50,54,55].

Another enzyme identified via explainable ML was acetyl-CoA carboxylase, which utilises a 2-step mechanism to carboxylate acetyl-CoA in fatty acid synthesis. The first step involves the ATP-dependent carboxylation of the cofactor biotin at its ureido ring. Several drugs depleted by the enzyme-encoding strain were found to bind the enzyme’s adenine-binding site with moderate affinity. Given the structural dissimilarities between these drugs and biotin, we propose that the drugs bind the enzyme but are not metabolised, instead forming a non-metabolised enzyme-drug complex.

Further work is needed to determine the optimal *k*-mer and/or NLP pipeline methodologies. Here, we investigated sequential *k*-mers and count vectorisation for tokenisation due to their low computational demands. Future work may instead explore sliding *k*-mers and other genomic featurisation techniques [56–60]. In the current methodology, it was found that strains were not equally represented by large *k*-mer sequences: several sequences in larger *k*-mers datasets were only found in the genome of one strain. What’s more, the present pipeline does not consider synonymous codon usage and instead treats functionally identical genetic sequences (like ‘TCAGGA’ and ‘TCAGGG’) as different sequences despite both encoding the serine-glycine amino acid sequence. Future investigations may therefore integrate understanding of DNA’s degenerate nature to create less-sparse datasets.

Alternate methods of drug representation, such as Mordred fingerprinting or those used by Sharma et al., may also be explored [61–63]. Additionally, metabolism of drug compounds does not always result in complete inactivation of drugs, and may lead to activation or toxification [64,65]. Thus, future models may incorporate information on metabolic enzyme activity to classify depletion instances into subclasses, shedding light on the biological impact of metabolism [61,62].

Bacterial drug depletion is a substantial concern, that remains neglected largely due to issues in *in vitro* studies. Billions of individuals suffer from gastrointestinal disorders, and close to the entire global population benefit from oral medications [66]. The investment of time, money, and resources into developing, manufacturing, distributing and prescribing these medications is questionable if they fail to tangibly benefit a considerable proportion of patients. Some drugs are depleted by all bacteria studied – the presence of a single strain can greatly diminish drug bioavailability and efficacy. This necessitates higher doses, raising costs, and the likelihood of adverse effects and toxicity [67].

Application of predictive models to novel drug compounds can provide pharmaceutical researchers with insights into which compounds are likely to be depleted in the average patient. Implementation of this screening in early stages of drug development may streamline the drug development pipeline, making the process more cost-effective and resource-efficient, while accelerating the transition from bench-to-bedside [68,69]. Insights on bacterial metabolism may also be applied beyond pharmaceutical sciences; for instance, in bioremediation efforts and the development of biodegradable plastics [70,71].

### Conclusions

Bacterial drug depletion is a significant concern for clinicians and the pharmaceutical industry but remains overlooked due to resource limitations and implementation challenges *in vitro* and *in vivo*. Research herein aims to systematically evaluate ML techniques and *k-*mer sizes in *in silico* drug depletion prediction, as well as provide mechanistic insights into depletion. This was achieved through a novel methodology; the integration of feature importance analysis, BLAST searching, and molecular docking simulations. ML models developed achieved markedly higher balanced accuracy scores than pre-existing models. F1 score found the best-performing ML model to be RF with 600-mers (0.59 ± 0.09). Feature importance analysis of this model genomic sequences vital for ML decision making. Subsequent BLAST annotation associated 2 candidate sequences with a glycoside hydrolase enzyme, and a a acetyl-CoA carboxylase biotin carboxylase subunit family protein. The glycoside hydrolase-encoding strain depleted all cardiac glycoside drugs investigated, a previously unreported phenomenon. It is proposed this enzyme cleaves the glycosidic bond between steroidal nucleus and saccharide ring. This is supported by molecular docking simulations revealing cardiac glycosides bind this enzyme with high binding affinity. Here, we demonstrate a novel methodology that accelerates mechanistic understanding of bacterial metabolism without *ex silico* studies. Within pharmaceutical settings, application of this research to identification of low bioavailability leads may drive down drug discovery costs, accelerate the transition from bench-to-bedside, and improve patient outcomes.

## Statements & Declarations

### Author Contributions

M.E. conceived and designed the study. N.F.A.K. performed the data collection. N.F.A.K. and M.E. contributed to the analysis, visualisation and first draft writing. All authors commented on previous versions of the manuscript. All authors read and approved the final manuscript.

### Data Availability

The datasets generated during and/or analysed during the current study are available from the corresponding author on reasonable request.

## Appendix

**Table S1:**
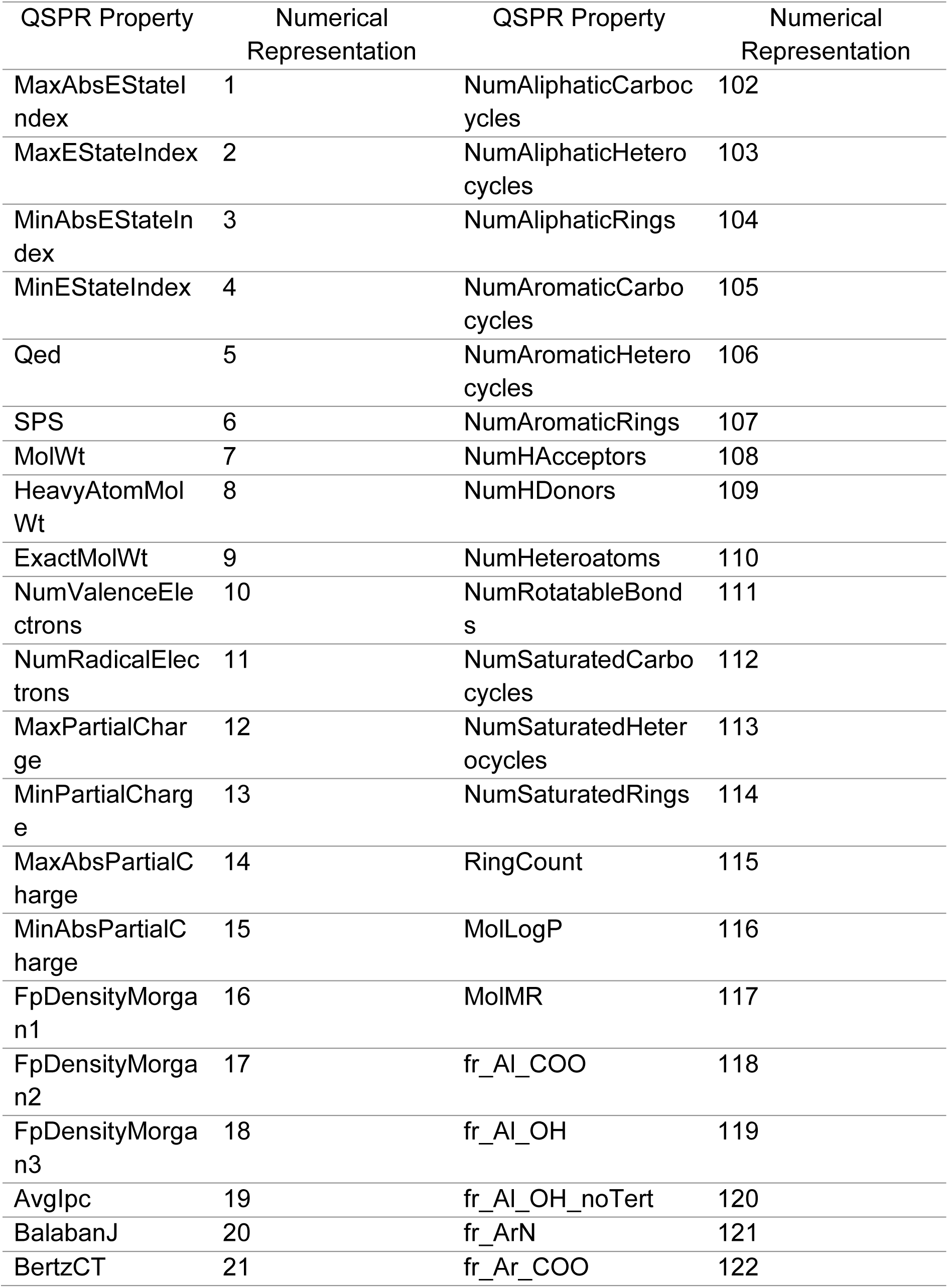
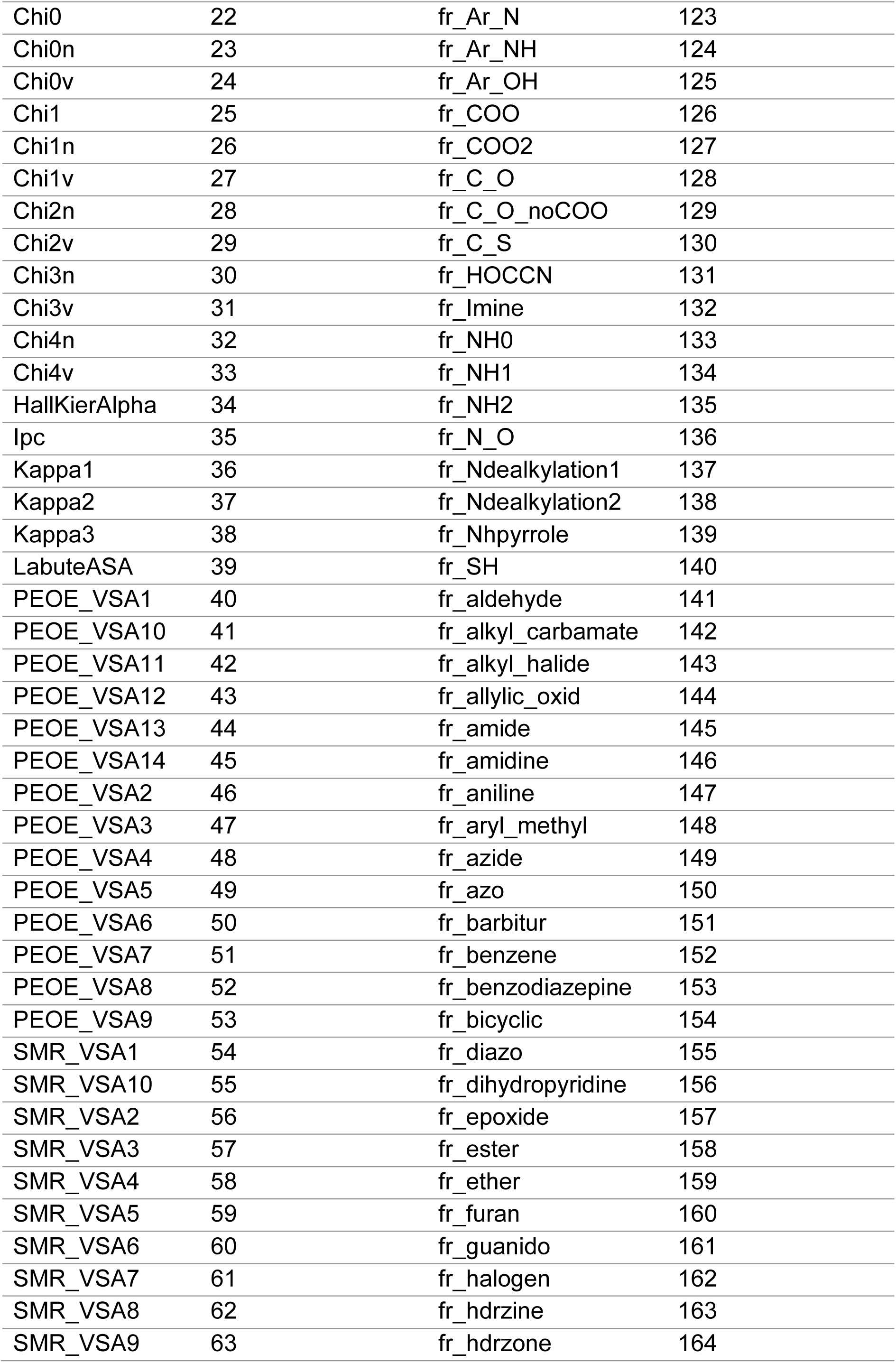
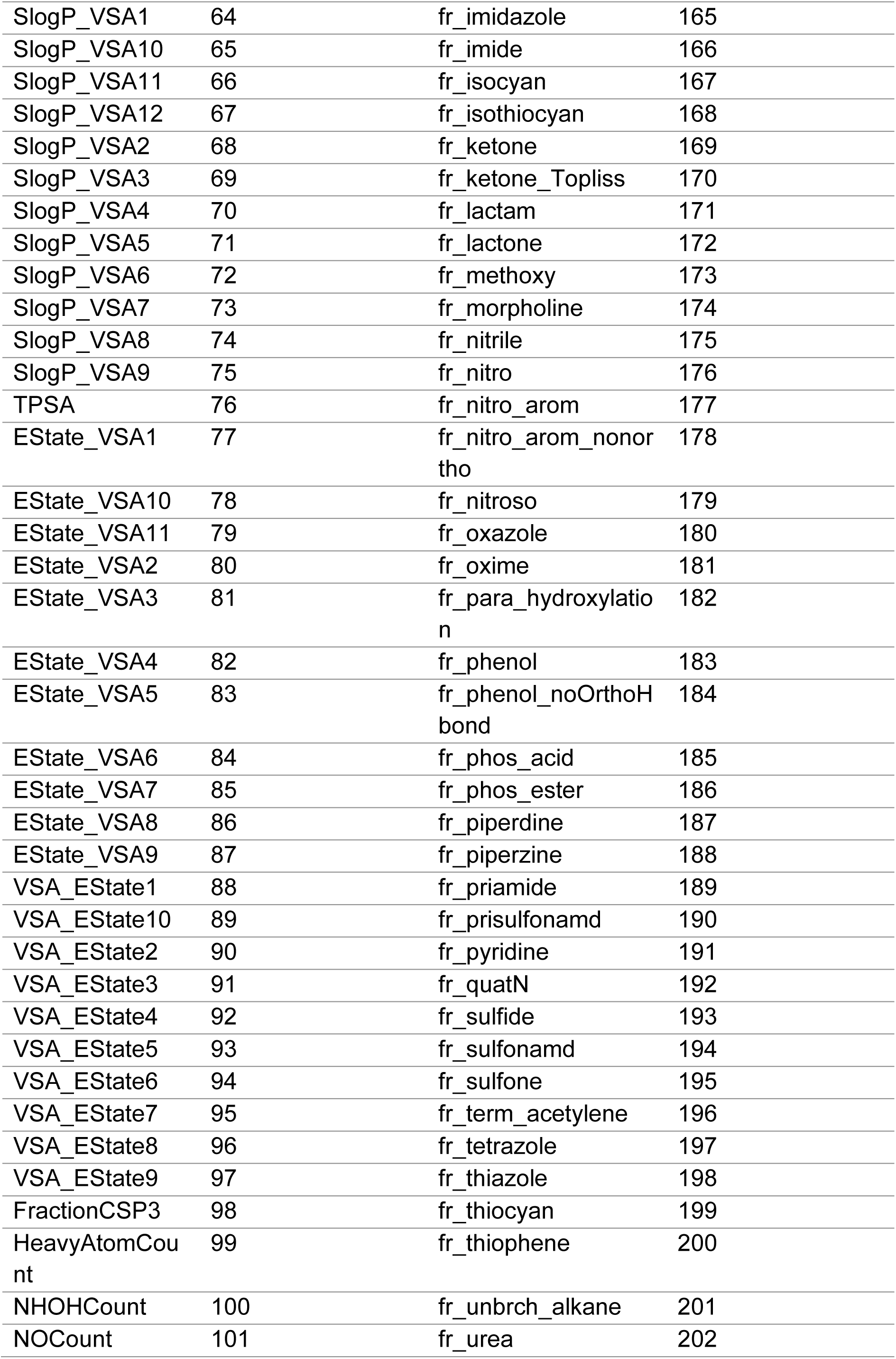
Numerical representations of QSPRs. . Refer to RDkit’s documentation for further information on structural properties.

## Reference List

[1] Alqahtani MS, Kazi M, Alsenaidy MA, Ahmad MZ. Advances in Oral Drug Delivery. Front Pharmacol 2021;12. 10.3389/fphar.2021.618411.

[2] Milián-Guimerá C, Tollemeto M, Mortensen JS, Badillo-Ramírez I, Jernskæg DR, Sam M, et al. Fabrication and characterization of enteric microneedle patches for oral delivery of small and macromolecule compounds. Int J Pharm 2026;687. 10.1016/j.ijpharm.2025.126376.

[3] Brown TD, Whitehead KA, Mitragotri S. Materials for oral delivery of proteins and peptides. Nat Rev Mater 2020;5:127–48. 10.1038/s41578-019-0156-6.

[4] Zhang X, Chen G, Zhang H, Shang L, Zhao Y. Bioinspired oral delivery devices. Nature Reviews Bioengineering 2023;1:208–25. 10.1038/s44222-022-00006-4.

[5] Liu L, McClements DJ, Liu X, Liu F. Overcoming Biopotency Barriers: Advanced Oral Delivery Strategies for Enhancing the Efficacy of Bioactive Food Ingredients. Advanced Science 2024;11. 10.1002/advs.202401172.

[6] Ejazi SA, Louisthelmy R, Maisel K. Mechanisms of Nanoparticle Transport across Intestinal Tissue: An Oral Delivery Perspective. ACS Nano 2023;17:13044–61. 10.1021/acsnano.3c02403.

[7] Elbadawi M, Nikjoo D, Gustafsson T, Gaisford S, Basit AW. Pressure-assisted microsyringe 3D printing of oral films based on pullulan and hydroxypropyl methylcellulose. Int J Pharm 2021;595. 10.1016/j.ijpharm.2021.120197.

[8] Zimmermann M, Zimmermann-Kogadeeva M, Wegmann R, Goodman AL. Mapping human microbiome drug metabolism by gut bacteria and their genes. Nature 2019;570:462–7. 10.1038/s41586-019-1291-3.

[9] McCoubrey LE, Elbadawi M, Basit AW. Current clinical translation of microbiome medicines. Trends Pharmacol Sci 2022;43:281–92. 10.1016/j.tips.2022.02.001.

[10] Zu M, Ma Y, Cannup B, Xie D, Jung Y, Zhang J, et al. Oral delivery of natural active small molecules by polymeric nanoparticles for the treatment of inflammatory bowel diseases. Adv Drug Deliv Rev 2021;176. 10.1016/j.addr.2021.113887.

[11] Heintz-Buschart A, Wilmes P. Human Gut Microbiome: Function Matters. Trends Microbiol 2018;26:563–74. 10.1016/j.tim.2017.11.002.

[12] Coombes Z, Yadav V, McCoubrey LE, Freire C, Basit AW, Conlan RS, et al. Progestogens are metabolized by the gut microbiota: Implications for colonic drug delivery. Pharmaceutics 2020;12:1–10. 10.3390/pharmaceutics12080760.

[13] He S, Li H, Yu Z, Zhang F, Liang S, Liu H, et al. The Gut Microbiome and Sex Hormone-Related Diseases. Front Microbiol 2021;12. 10.3389/fmicb.2021.711137.

[14] Vinarov Z, Abdallah M, Agundez JAG, Allegaert K, Basit AW, Braeckmans M, et al. Impact of gastrointestinal tract variability on oral drug absorption and pharmacokinetics: An UNGAP review. European Journal of Pharmaceutical Sciences 2021;162. 10.1016/j.ejps.2021.105812.

[15] Klünemann M, Andrejev S, Blasche S, Mateus A, Phapale P, Devendran S, et al. Bioaccumulation of therapeutic drugs by human gut bacteria. Nature 2021;597:533–8. 10.1038/s41586-021-03891-8.

[16] Javdan B, Lopez JG, Chankhamjon P, Lee YCJ, Hull R, Wu Q, et al. Personalized Mapping of Drug Metabolism by the Human Gut Microbiome. Cell 2020;181:1661–1679.e22. 10.1016/j.cell.2020.05.001.

[17] McCoubrey LE, Gaisford S, Orlu M, Basit AW. Predicting drug-microbiome interactions with machine learning. Biotechnol Adv 2022;54. 10.1016/j.biotechadv.2021.107797.

[18] Awad A, Madla CM, McCoubrey LE, Ferraro F, Gavins FKH, Buanz A, et al. Clinical translation of advanced colonic drug delivery technologies. Adv Drug Deliv Rev 2022;181. 10.1016/j.addr.2021.114076.

[19] Richter MF, Drown BS, Riley AP, Garcia A, Shirai T, Svec RL, et al. Predictive compound accumulation rules yield a broad-spectrum antibiotic. Nature 2017;545:299–304. 10.1038/nature22308.

[20] McCoubrey LE, Basit AW. Addressing drug-microbiome interactions: the role of healthcare professionals. Pharm J 2022;308:1–27. 10.1211/PJ.2022.1.126780.

[21] Lee JR, Muthukumar T, Dadhania D, Taur Y, Jenq RR, Toussaint NC, et al. Gut microbiota and tacrolimus dosing in kidney transplantation. PLoS One 2015;10. 10.1371/journal.pone.0122399.

[22] van Kessel SP, Frye AK, El-Gendy AO, Castejon M, Keshavarzian A, van Dijk G, et al. Gut bacterial tyrosine decarboxylases restrict levels of levodopa in the treatment of Parkinson’s disease. Nat Commun 2019;10. 10.1038/s41467-019-08294-y.

[23] Yadav V, Mai Y, Mccoubrey LE, Wada Y, Tomioka M, Kawata S, et al. 5-Aminolevulinic Acid as a Novel Therapeutic for Inflammatory Bowel Disease. Biomedicines 2021;9. 10.3390/biomedicines.

[24] Yadav V, Varum F, Bravo R, Furrer E, Basit AW. Gastrointestinal stability of therapeutic anti-TNF α IgG1 monoclonal antibodies. Int J Pharm 2016;502:181–7. 10.1016/j.ijpharm.2016.02.014.

[25] Tannergren C, Borde A, Boreström C, Abrahamsson B, Lindahl A. Evaluation of an in vitro faecal degradation method for early assessment of the impact of colonic degradation on colonic absorption in humans. European Journal of Pharmaceutical Sciences 2014;57:200–6. 10.1016/j.ejps.2013.10.001.

[26] McCoubrey LE, Elbadawi M, Orlu M, Gaisford S, Basit AW. Machine learning uncovers adverse drug effects on intestinal bacteria. Pharmaceutics 2021;13. 10.3390/pharmaceutics13071026.

[27] McCoubrey LE, Seegobin N, Elbadawi M, Hu Y, Orlu M, Gaisford S, et al. Active Machine learning for formulation of precision probiotics. Int J Pharm 2022;616. 10.1016/j.ijpharm.2022.121568.

[28] McCoubrey LE, Thomaidou S, Elbadawi M, Gaisford S, Basit AW, Orlu M. Machine learning predicts drug metabolism and bioaccumulation by intestinal microbiota. Pharmaceutics 2021;13. 10.3390/pharmaceutics13122001.

[29] Mallory EK, Acharya A, Rensi SE, Turnbaugh PJ, Bright RA, Altman RB. Chemical reaction vector embeddings: towards predicting drug metabolism in the human gut microbiome. vol. 23. 2018.

[30] Elmassry MM, Kim S, Busby B. Predicting drug-metagenome interactions: Variation in the microbial β-glucuronidase level in the human gut metagenomes. PLoS One 2021;16. 10.1371/journal.pone.0244876.

[31] Elbadawi M, Gaisford S, Basit AW. Advanced machine-learning techniques in drug discovery. Drug Discov Today 2021;26:769–77. 10.1016/j.drudis.2020.12.003.

[32] Reker D, Shi Y, Kirtane AR, Hess K, Zhong GJ, Crane E, et al. Machine Learning Uncovers Food- and Excipient-Drug Interactions. Cell Rep 2020;30:3710–3716.e4. 10.1016/j.celrep.2020.02.094.

[33] Gavins FKH, Fu Z, Elbadawi M, Basit AW, Rodrigues MRD, Orlu M. Machine learning predicts the effect of food on orally administered medicines. Int J Pharm 2022;611. 10.1016/j.ijpharm.2021.121329.

[34] Wang F, Sangfuang N, McCoubrey LE, Yadav V, Elbadawi M, Orlu M, et al. Advancing oral delivery of biologics: Machine learning predicts peptide stability in the gastrointestinal tract. Int J Pharm 2023;634. 10.1016/j.ijpharm.2023.122643.

[35] Azodi CB, Tang J, Shiu SH. Opening the Black Box: Interpretable Machine Learning for Geneticists. Trends in Genetics 2020;36:442–55. 10.1016/j.tig.2020.03.005.

[36] O’Leary NA, Cox E, Holmes JB, Anderson WR, Falk R, Hem V, et al. Exploring and retrieving sequence and metadata for species across the tree of life with NCBI Datasets. Sci Data 2024;11. 10.1038/s41597-024-03571-y.

[37] García-López M, Meier-Kolthoff JP, Tindall BJ, Gronow S, Woyke T, Kyrpides NC, et al. Analysis of 1,000 Type-Strain Genomes Improves Taxonomic Classification of Bacteroidetes. Front Microbiol 2019;10. 10.3389/fmicb.2019.02083.

[38] Madden T. The BLAST Sequence Analysis Tool. 2013.

[39] Ali I, Khan A, Fa Z, Khan T, Wei DQ, Zheng J. Crystal structure of Acetyl-CoA carboxylase (AccB) from Streptomyces antibioticus and insights into the substrate-binding through in silico mutagenesis and biophysical investigations. Comput Biol Med 2022;145. 10.1016/j.compbiomed.2022.105439.

[40] Mochalkin I, Miller JR, Narasimhan L, Thanabal V, Erdman P, Cox PB, et al. Discovery of antibacterial biotin carboxylase inhibitors by virtual screening and fragment-based approaches. ACS Chem Biol 2009;4:473–83. 10.1021/cb9000102.

[41] Cifone MT, He Y, Basu R, Wang N, Davoodi S, Spagnuolo LA, et al. Heterobivalent Inhibitors of Acetyl-CoA Carboxylase: Drug Target Residence Time and Time-Dependent Antibacterial Activity. J Med Chem 2022;65:16510–25. 10.1021/acs.jmedchem.2c01380.

[42] Broussard TC, Pakhomova S, Neau DB, Bonnot R, Waldrop GL. Structural Analysis of Substrate, Reaction Intermediate, and Product Binding in Haemophilus influenzae Biotin Carboxylase. Biochemistry 2015;54:3860–70. 10.1021/acs.biochem.5b00340.

[43] Mochalkin I, Miller JR, Evdokimov A, Lightle S, Yan C, Stover CK, et al. Structural evidence for substrate-induced synergism and half-sites reactivity in biotin carboxylase. Protein Science 2008;17:1706–18. 10.1110/ps.035584.108.

[44] Elbadawi M, McCoubrey LE, Gavins FKH, Ong JJ, Goyanes A, Gaisford S, et al. Disrupting 3D printing of medicines with machine learning. Trends Pharmacol Sci 2021;42:745–57. 10.1016/j.tips.2021.06.002.

[45] Uddin S, Lu H. Confirming the statistically significant superiority of tree-based machine learning algorithms over their counterparts for tabular data. PLoS One 2024;19. 10.1371/journal.pone.0301541.

[46] Kumar R, Henrissat B, Coutinho PM. Intrinsic dynamic behavior of enzyme:substrate complexes govern the catalytic action of β-galactosidases across clan GH-A. Sci Rep 2019;9. 10.1038/s41598-019-46589-8.

[47] Ioannou A, Knol J, Belzer C. Microbial Glycoside Hydrolases in the First Year of Life: An Analysis Review on Their Presence and Importance in Infant Gut. Front Microbiol 2021;12. 10.3389/fmicb.2021.631282.

[48] Liu X, Li J, Wu R, Bai L. Characterization of glycogen-related glycoside hydrolase glgX and glgB from Klebsiella pneumoniae and their roles in biofilm formation and virulence. Front Cell Infect Microbiol 2024;14. 10.3389/fcimb.2024.1507332.

[49] Haiser HJ, Gootenberg DB, Chatman K, Sirasani G, Balskus EP, Turnbaugh PJ. Predicting and manipulating cardiac drug inactivation by the human gut bacterium Eggerthella lenta. Science (1979) 2013;341:295–8. 10.1126/science.1235872.

[50] Ricaurte D, Huang Y, Sheth RU, Gelsinger DR, Kaufman A, Wang HH. High-throughput transcriptomics of 409 bacteria–drug pairs reveals drivers of gut microbiota perturbation. Nat Microbiol 2024;9:561–75. 10.1038/s41564-023-01581-x.

[51] Sinha AK, Laursen MF, Brinck JE, Rybtke ML, Hjørne AP, Procházková N, et al. Dietary fibre directs microbial tryptophan metabolism via metabolic interactions in the gut microbiota. Nat Microbiol 2024;9:1964–78. 10.1038/s41564-024-01737-3.

[52] Sheridan PO, Louis P, Tsompanidou E, Shaw S, Harmsen HJ, Duncan SH, et al. Distribution, organization and expression of genes concerned with anaerobic lactate utilization in human intestinal bacteria. Microb Genom 2022;8. 10.1099/mgen.0.000739.

[53] Arzamasov AA, Nakajima A, Sakanaka M, Ojima MN, Katayama T, Rodionov DA, et al. Human Milk Oligosaccharide Utilization in Intestinal Bifidobacteria Is Governed by Global Transcriptional Regulator NagR. MSystems 2022;7. 10.1128/msystems.00343-22.

[54] Culley C, Vijayakumar S, Zampieri G, Angione C. A mechanism-aware and multiomic machine-learning pipeline characterizes yeast cell growth. SYSTEMS BIOLOGY BIOPHYSICS AND COMPUTATIONAL BIOLOGY 2002;117:18869–79. 10.1073/pnas.2002959117/-/DCSupplemental.y.

[55] Smith LA, Cahill JA, Lee J-H, Graim K. Equitable machine learning counteracts ancestral bias in precision medicine. Nat Commun 2025;16:2144. 10.1038/s41467-025-57216-8.

[56] Ghandi M, Lee D, Mohammad-Noori M, Beer MA. Enhanced Regulatory Sequence Prediction Using Gapped k-mer Features. PLoS Comput Biol 2014;10. 10.1371/journal.pcbi.1003711.

[57] Ghandi M, Mohammad-Noori M, Beer MA. Robust k-mer frequency estimation using gapped k-mers. J Math Biol 2014;69:469–500. 10.1007/s00285-013-0705-3.

[58] Ji Y, Zhou Z, Liu H, Davuluri R V. DNABERT: pre-trained Bidirectional Encoder Representations from Transformers model for DNA-language in genome. Bioinformatics 2021;37:2112–20. 10.1093/bioinformatics/btab083.

[59] Koblitz J, Reimer LC, Pukall R, Overmann J. Predicting bacterial phenotypic traits through improved machine learning using high-quality, curated datasets. Commun Biol 2025;8. 10.1038/s42003-025-08313-3.

[60] Harling-Lee JD, Gorzynski J, Yebra G, Angus T, Fitzgerald JR, Freeman TC. A graph-based approach for the visualisation and analysis of bacterial pangenomes. BMC Bioinformatics 2022;23. 10.1186/s12859-022-04898-2.

[61] Sharma AK, Jaiswal SK, Chaudhary N, Sharma VK. A novel approach for the prediction of species-specific biotransformation of xenobiotic/drug molecules by the human gut microbiota. Sci Rep 2017;7. 10.1038/s41598-017-10203-6.

[62] Malwe AS, Srivastava GN, Sharma VK. GutBug: A Tool for Prediction of Human Gut Bacteria Mediated Biotransformation of Biotic and Xenobiotic Molecules Using Machine Learning. J Mol Biol 2023;435. 10.1016/j.jmb.2023.168056.

[63] Stubbs CD, Kim Y, Quinn EC, Pérez-Soto R, Chen EYX, Kim S. Predicting homopolymer and copolymer solubility through machine learning. Digital Discovery 2024. 10.1039/d4dd00290c.

[64] Koppel N, Rekdal VM, Balskus EP. Chemical transformation of xenobiotics by the human gut microbiota. Science (1979) 2017;356:1246–57. 10.1126/science.aag2770.

[65] Zimmermann M, Zimmermann-Kogadeeva M, Wegmann R, Goodman AL. Separating host and microbiome contributions to drug pharmacokinetics and toxicity. Science (1979) 2019;363. 10.1126/science.aat9931.

[66] Sperber AD, Bangdiwala SI, Drossman DA, Ghoshal UC, Simren M, Tack J, et al. Worldwide Prevalence and Burden of Functional Gastrointestinal Disorders, Results of Rome Foundation Global Study. Gastroenterology 2021;160:99–114.e3. 10.1053/j.gastro.2020.04.014.

[67] Hill AM, Barber MJ, Gotham D. Estimated costs of production and potential prices for the WHO essential medicines list. BMJ Glob Health 2018;3. 10.1136/bmjgh-2017-000571.

[68] Burt T, Button KS, Thom HHZ, Noveck RJ, Munafò MR. The Burden of the “False-Negatives” in Clinical Development: Analyses of Current and Alternative Scenarios and Corrective Measures. Clin Transl Sci 2017;10:470–9. 10.1111/cts.12478.

[69] Jithesh PV, Abuhaliqa M, Syed N, Ahmed I, El Anbari M, Bastaki K, et al. A population study of clinically actionable genetic variation affecting drug response from the Middle East. NPJ Genom Med 2022;7. 10.1038/s41525-022-00281-5.

[70] Cai Z, Li M, Zhu Z, Wang X, Huang Y, Li T, et al. Biological Degradation of Plastics and Microplastics: A Recent Perspective on Associated Mechanisms and Influencing Factors. Microorganisms 2023;11. 10.3390/microorganisms11071661.

[71] Omura T, Isobe N, Miura T, Ishii S, Mori M, Ishitani Y, et al. Microbial decomposition of biodegradable plastics on the deep-sea floor. Nat Commun 2024;15. 10.1038/s41467-023-44368-8.

